# DeepOM: Single-molecule optical genome mapping via deep learning

**DOI:** 10.1101/2022.11.04.512597

**Authors:** Yevgeni Nogin, Tahir Detinis Zur, Sapir Margalit, Ilana Barzilai, Onit Alalouf, Yuval Ebenstein, Yoav Shechtman

## Abstract

Efficient tapping into genomic information from a single microscopic image of an intact DNA molecule fragment is an outstanding challenge and its solution will open new frontiers in molecular diagnostics. Here, a new computational method for optical genome mapping utilizing Deep Learning is presented, termed DeepOM. Utilization of a Convolutional Neural Network (CNN), trained on simulated images of labeled DNA molecules, improves the success rate in alignment of DNA images to genomic references. The method is evaluated on acquired images of human DNA molecules stretched in nano-channels. The accuracy of the method is benchmarked against state-of-the-art commercial software Bionano Solve. The results show a significant advantage in alignment success rate for molecules shorter than 50 kb. DeepOM improves yield, sensitivity and throughput of optical genome mapping experiments in applications of human genomics and microbiology.

## 1 Introduction

Optical genome mapping (OGM) of DNA [1, 2, 3] involves the imaging of labeled DNA molecules and their alignment to reference genome sequences. Consequently, the resulting best-matching alignment reports on the exact position of this molecule fragment in one of the organism’s chromosomes. This information enables multiple applications in molecular diagnostics and in genomic research.

Example applications of OGM include species identification [4, 5, 6, 7] for applications such as pathogen identification in clinical samples, as well as genome-wide mapping of effects such as: DNA damage [8], methylation [9], and structural variations [10]. OGM holds several advantages compared to DNA sequencing; for one, it produces extremely long reads of potentially megabase size, which are necessary for mapping large-scale structural and copy number variations in the genome. Additionally, as a single-molecule technique, it holds the potential for extremely high sensitivity, i.e. detection of low quantities of target DNA [11], which is necessary in applications such as cultivation-free pathogen identification [7].

Given an image of a DNA molecule labeled at a specific sequence motif, multiple computational approaches have been proposed for its alignment to a reference genome sequence. If the labelling is sparse enough so that individual fluorescent labels can be separated, the positions of the labels are determined using standard localization techniques, such as emitter centroid fitting [12]. Then, Dynamic Programming algorithms [13] are employed to align the label positions to the expected positions of the labeled motif in a reference genome sequence. When the labeled motif is dense in the genome and does not allow for separation of individual labels, a different approach was used [4, 5, 6, 7], in which the intensity profile along the imaged molecule is aligned by cross-correlation to the theoretical intensity profile expected from the density of the labeled motif in the reference genome.

The accuracy of OGM can be defined as the expected fraction of imaged molecules that are aligned with high confidence to the reference genome. This accuracy is extremely important for applications where target DNA quantity in the sample is limited, such as cultivation-free pathogen identification [7], or where maximal coverage of the genome is required per mapping experiment, such as in rare variant detection [14] or epigenetic mapping [15, 16]. The current computational approaches are limited in accuracy, since they are unable to extract all the available information from the image of the DNA molecule. Specifically, when emitters are overlapping inside a diffraction-limited spot, classic approaches usually cannot separate them.

In this study, in order to maximise the information extracted from the molecule image, a Deep Learning approach is presented. Convolutional Neural Networks (CNN) were previously shown to become the state-of-the-art for Single Molecule Localization Microscopy (SMLM) [17, 18, 19]. Here, a similar approach is applied to OGM, and its advantage is demonstrated on images of sparsely-labeled DNA molecules stretched in nanochannels. The alignment accuracy of the presented method DeepOM is compared against the commercial Bionano Solve software which localizes sparse emitters, neglecting their diffraction-limited image overlap. In contrast, the localization neural network of DeepOM enables the separation of multiple fluorescent emitters that are within a diffraction limited spot. Since the probability for wrong alignment of optical maps, as was theoretically shown, depends exponentially on the number of localized labels in the query molecule [20], the detection of more labels per kilobase of DNA by DeepOM, results in a significantly higher alignment success rate.

## 2 Materials and Methods

### 2.1 The DeepOM Method

The DeepOM alignment of a DNA molecule to a reference genome sequence starts from query images of molecules fluorescently labeled at specific motifs (Figure 1). The motif CTTAAG (referred to as DLE-1 by Bionano Genomics) was labeled in this study. In each molecule image, the labels are localized by a localization neural network, resulting in a query map of 1-D pixel positions of labels along the length of the molecule. A reference map is the sequence of base-pair positions of the labeled motif in a reference genome sequence. The resulting query map is aligned to the reference map with the dynamic programming alignment algorithm presented below.

**Figure 1:**
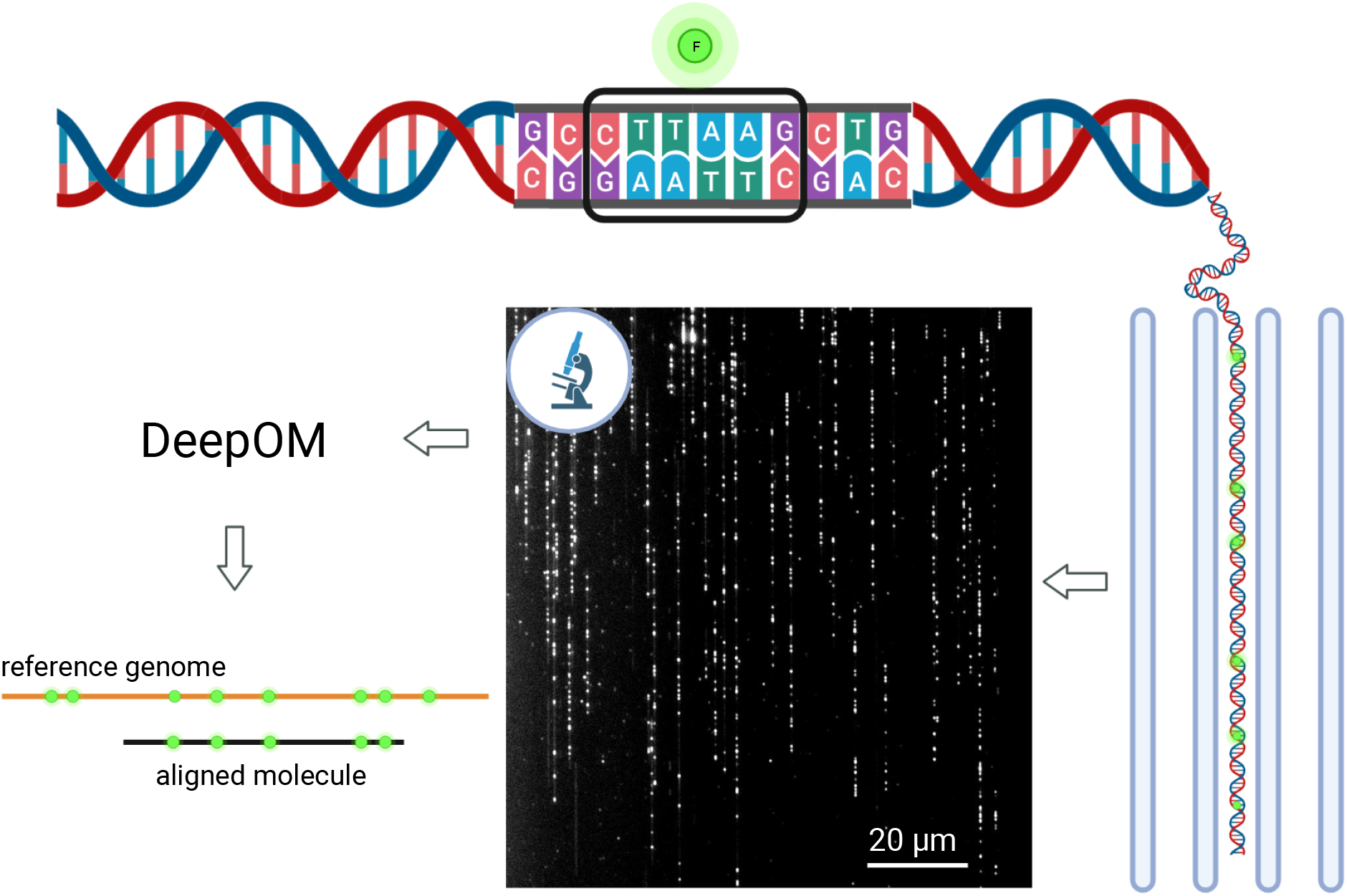
Optical genome mapping using DeepOM. DNA molecules are fluorescently labeled at specific sequence motifs, CTTAAG in this study. Then, they are stretched in nano-channels and imaged in a microscope. The images are analyzed by the DeepOM software, and each molecule is aligned to its top matching position in one of the reference genome sequences.

#### Localization Neural Network

A localization neural net model was trained following DeepStorm [17], DeepStorm3D [18], and DECODE [19], where the models are trained in a supervised manner on simulated images from randomly generated ground-truth emitter positions, derived using an optical forward model. Here, a 2-D Gaussian point-spread function (PSF) was used for the optical forward model, and emitter positions were confined to a straight line segment (Figure 2). Following DECODE [19], and DeepStorm [17], a U-Net [21] was used, but with 1-D convolutional layers instead of 2-D convolutional layers. The input image to the network, which is usually on the order of 5 pixels wide and an order of 100 pixels long, was regarded as a 1-D image with the width (or lateral) dimension regarded as neural network channels dimension. The last layer of the U-Net was modified to output two 1-D vectors, which correspond to two output numbers per 1-D pixel: (a) Occupancy probability, i.e. the probability for having an emitter in a pixel, and (b) relative position of the emitter inside the pixel, if the pixel contains an emitter. This is valid assuming there is at most one emitter per pixel, which is a good approximation for most datasets of interest, including the one presented here. For the two neural network output numbers defined above, the loss *ℒ* is computed as a sum of two loss terms: the occupancy loss *ℒ*_*occ*_, and the localization loss *ℒ*_*loc*_,

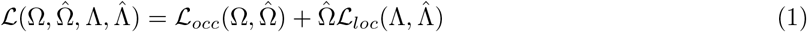

where Ω_*i*_ is the predicted probability for having an emitter in a pixel *i*; 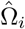 is the ground-truth emitter existence in the pixel, equal to 1 if an emitter is in a pixel and 0 otherwise; Λ_*i*_ is the relative position of an emitter inside the pixel ranging from 0 for the left pixel edge to 1 for the right pixel edge. This position has a meaningful value only in the pixels containing emitters, so the localization loss *ℒ*_*loc*_ is masked with 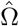 in the equation; 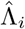 is the ground-truth relative position of an emitter inside the pixel computed from ground-truth emitter positions, which are known in the simulated data. The loss terms themselves were computed as,

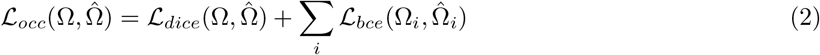

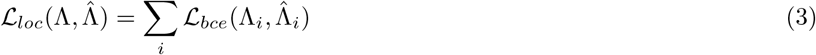

where the *i*-summation is over the 1-D pixel indices along the length of the molecule image, and with Dice-Loss [22] *ℒ*_*dice*_ and Binary-Cross-Entropy *ℒ*_*bce*_ defined as,

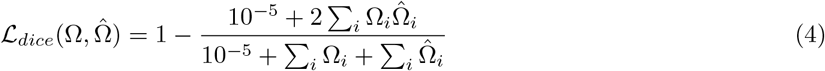

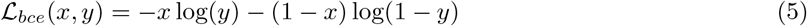

In each gradient descent training step, the model was presented with a batch of randomly generated DNA molecule images, and the loss was computed as described above using the ground truth positions of emitters in the molecule (Figure 2). Training was done for 10000 steps with a 256 batch size.

**Figure 2:**
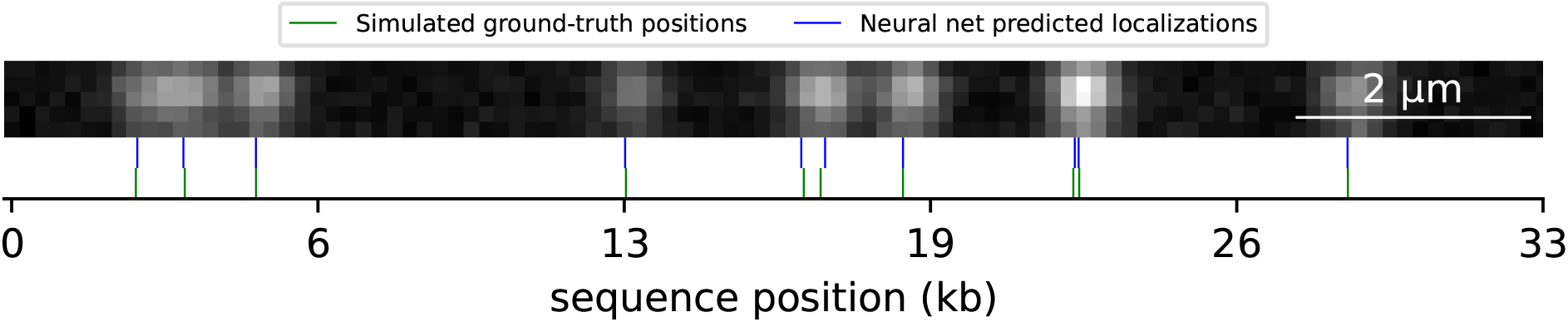
Simulated DNA molecule used for localizer neural net training. Ground-truth label positions (green) and the predicted localizations by the neural net (blue) are shown. Random generated emitter positions were confined to a straight line segment, convolved with a 2-D Gaussian point-spread function (PSF), and noise was added to the image.

#### Alignment algorithm

The algorithm by Valouev et al. [13] was implemented to align the localized labels in a DNA molecule to the reference genome. An alignment of a DNA query molecule and a reference genome sequence is a set of labeled position pairs from the query and reference. The implementation of the algorithm computes the following Dynamic Programming recurrence equation for the alignment score matrix *S*_*i,j*_,

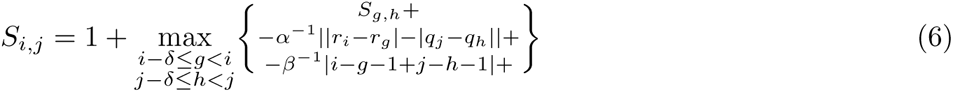

Then, the alignment is traversed back starting from the maximal cell value in the score matrix. *r*_*i*_ is the reference positions vector indexed by the integers *i* or *g*, and *q*_*j*_ is the query positions vector indexed by the integers *j* or *h*. The query vector *q* = *sx* is obtained by converting the pixel value *x* of localized emitters to basepairs through a conversion scale factor 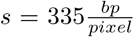. *δ* = 5 is the allowed margin for missing labels in query or reference, *α* = 500 is the penalty factor for localization error, *β* = 10 is the penalty factor for a missing label in the alignment. *S*_*i,j*_ is the score of the top scoring alignment of *r*_*i*_ and *q*_*j*_ ending in indices *i, j*.

### 2.2 Sample preparation and imaging

#### Cell culture

U2OS (human OS) cell line was cultured in Dulbecco’s Modified Eagle medium, supplemented with 10% fetal bovine serum (Gibco, Amarillo, TX), 2 mM l-glutamine, and 1% penicillin-streptomycin (10,000 U/mL; Gibco) and incubated at 37°C with 5% CO2.

#### DNA extraction

DNA was extracted from 10^6^ cells using the Bionano Prep Cell Culture DNA Isolation Protocol according to manufacturer’s instructions.

#### DNA labeling

1 µg of DNA was directly labeled and stained using DLS labeling kit (Bionano Genomics) composed of a single enzymatic labeling reaction with DLE-1 enzyme followed by DNA staining with a fluorescent marker. 1 µg of DNA was mixed with 6 µl of 5x DLE-buffer, 2 µl of 20x DL-Green and 2 µl of DLE-1 enzyme (Bionano Genomics) in a total reaction volume of 30 µl and incubated for 2 hours at 37°C.

#### DNA imaging

DNA image data was generated on the Saphyr instrument (Bionano Genomics) with Saphyr chips (G1.2). The chip was loaded as recommended by Bionano Genomics.

## 3 Results and Discussion

The accuracy of DeepOM’s alignments was evaluated on images (Figure 3) produced from the Bionano Genomics Saphyr system described in the Materials and Methods section. The reference genome used for alignments is the reference human genome GRCh38 [24]. The ground-truth for alignments was generated as follows. All molecules longer than 450 kb were taken from the imaged data, and aligned to the reference genome with the Bionano Solve software. Out of those, the top 512 molecules were chosen by their Bionano alignment confidence score (see Bionano documentation [25]). Each chosen long molecule image was digitally cropped [26] into random short fragments (Figure 3). Since each cropped fragment’s position is known within the parent molecule, its aligned position can be regarded as a ground-truth for the purpose of the alignment accuracy evaluation. Each cropped fragment image is fed into the DeepOM pipeline and aligned to the reference genome, then if the alignment matches the ground-truth it is counted as correct.

**Figure 3:**
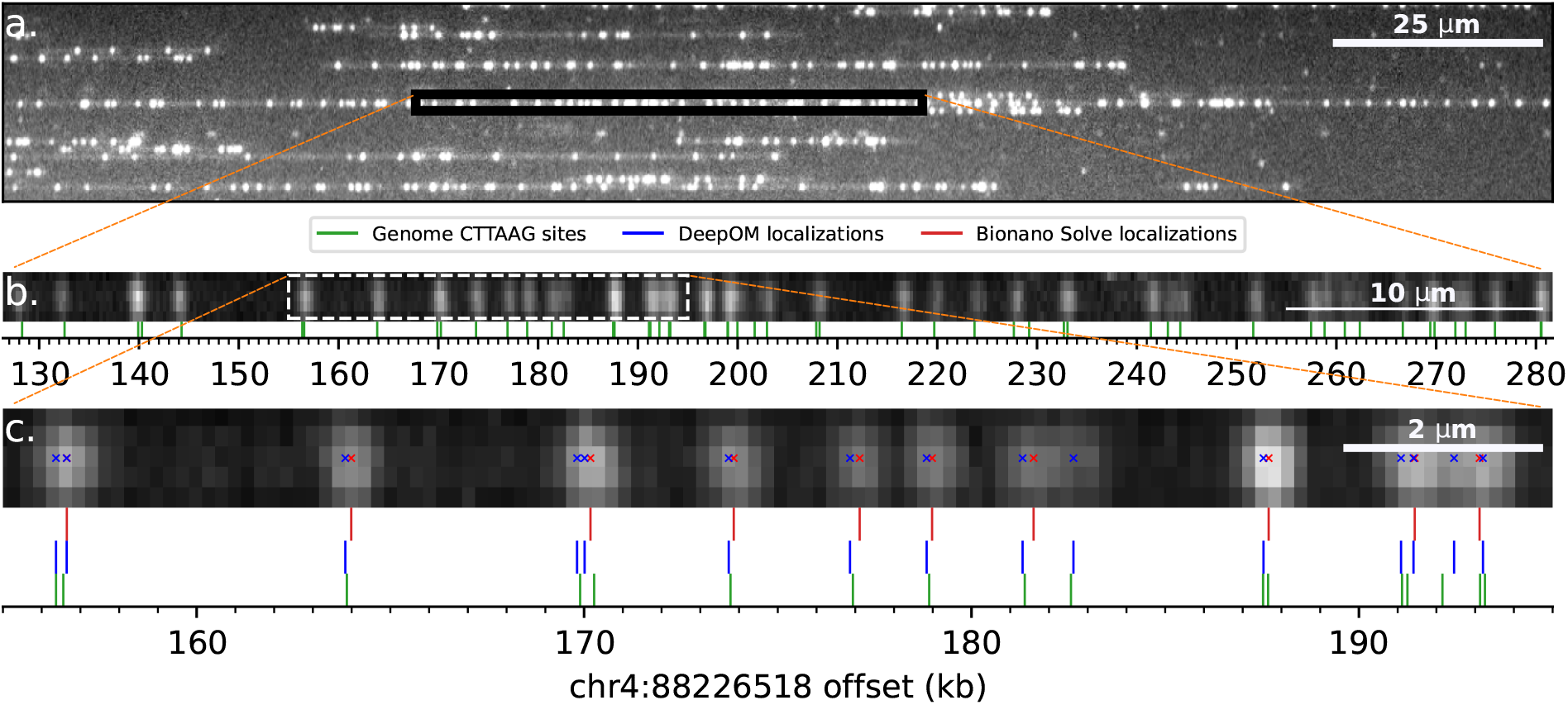
Experimentally imaged DNA molecules and their alignment to the human genome. **(a)** Zoomed-in field-of-view of an image captured in the Bionano Saphyr system. DNA molecules are stretched here in nano-channels of the Bionano Chip. **(b)** Zoomed-in view to a 500kb molecule from the field-of-view image (sub-figure a). This molecule is used as a ground-truth for alignment of its cropped sub-fragments. Cropped sub-fragment (white dashed rectangle), zoomed-in in sub-figure c. The reference genome labeled motif (CTTAAG) sites are shown (green), in relative offset to human genome coordinates shown on the x-axis. The alignment of the molecule to the reference was done both by Bionano Solve and DeepOM, and the resulting genome coordinates were practically identical. **(c)** An example cropped fragment used for the alignment success rate comparison. Shown are Bionano Solve localizations (red), DeepOM localizations (blue). The reference sites (green) genome coordinates of the parent molecule are used as a ground-truth for the evaluation of the success of this fragment’s alignment. The advantage of DeepOM is manifested here, where pairs of tightly spaced labeled motifs are separated by the neural net, while the classical localization approach (red) detects only one label at the diffraction limited spot. This in turn, leads to more confident and accurate alignments with higher success rates.

To make the comparison to the Bionano Solve software, cropping of the long molecules was done digitally by manipulation of the Bionano BNX output files (see documentation [25]) produced from the imaging experiment. The BNX file contains a localization list for the emitters in each molecule, and the molecules’ coordinates in the captured field-of-view image. In order to generate the cropped fragments in the BNX file, labels were deleted from the localization list according to the cropped fragments (Figure 3). First, we demonstrate the effectiveness of dense localization by the neural net, compared to standard, sparse localization, which discards closely spaced emitters. To do so, the localization list for each cropped molecule was aligned to the reference using the DeepOM alignment algorithm (Materials and Methods), and the success rate is shown in Figure 4a, presenting the comparison of the DeepOM localizer and the Bionano localizer both using the same alignment algorithm of DeepOM.

**Figure 4:**
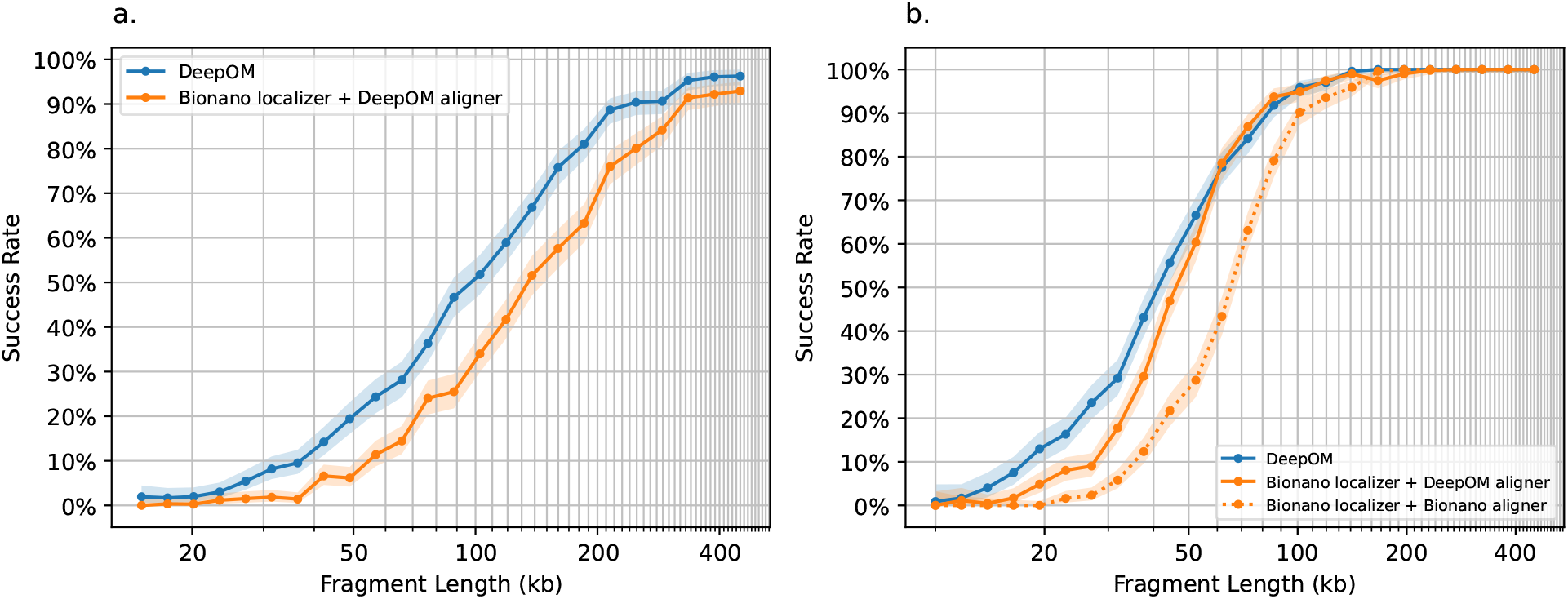
Accuracy evaluation of DeepOM vs Bionano Solve. The alignment success rate is shown for DeepOM (blue) and Bionano Solve (orange) vs fragment length. The success rate for a fragment length, is defined as the fraction of correct alignments for this length. Each success rate point was computed from 512 random cropped fragments with a ground-truth alignment from long high-confidence aligned molecules (Figure 3). Each cropped fragment was aligned to the reference genome and the success rate is the fraction of correct alignments, 95% confidence bounds are shown, computed with the Clopper-Pearson interval Beta Distribution [23]. In **(a)** DeepOM is compared against the localizations produced by Bionano Solve, which are aligned to the reference genome, with the DeepOM aligner. While in **(b)** comparison is also against the whole Bionano Solve pipeline including the Bionano localizer and Bionano aligner (orange dotted line).

Next, the whole DeepOM pipeline was compared vs the full Bionano localizer and aligner pipeline, on a subset of molecules. To run the Bionano pipeline, a new BNX input file was generated containing random crops from the chosen molecules. Then the file was fed as input to the Bionano Solve aligner. The success rate comparison of the full pipelines is shown in Figure 4b.

In both comparison methods, the results in Figure 4 show more than a twofold improvement factor in the success rate for fragments shorter than 50 kb. Notably, optimizing the alignment algorithm, together with the localization neural net, further improves alignment success rate, as can be seen when comparing the DeepOM aligner to the Bionano aligner, when both using the same localizer (Figure 4b).

## 4 Conclusions

In this study, an improved computational method for optical genome mapping was presented. A CNN was employed to significantly improve the success rate of alignments, as compared to a state-of-the-art non-overlapping approach. The accuracy of the presented method, DeepOM, was compared against the state-of-the-art commercial Bionano Solve on human cell-line DNA data acquired with the Bionano Saphyr system. The advantage of the presented method is most dominant for DNA fragments in the range 50-150kb, where it yields up to twofold more successful alignments (Figure 4b). This is especially significant given that the Bionano Genomics pipeline recommends filtering out molecules shorter than 150kb in order to provide a high mapping rate. In contast, DeepOM allows exploiting the information from these shorter molecules. DeepOM enables higher genome coverage from a given sample, enhancing the ability to detect low frequency structural variations. The presented method may be utilized in molecular diagnostic applications such as epigenetic profiling, and pathogen species identification, where it can significantly increase the fraction of identified molecules, enabling higher diagnostic sensitivity.

## 5 Acknowledgements

We thank Daniella Bar-Lev, Dganit Hanania-Buchris, Itai Orr, Eitan Yaakobi, Johan Hofkens, Robert Neely, Elias Nehme, Guy Tennenholtz, Jonathan Jeffet, Thomas Dages, Antoine Vinciguerra, Roy Velich, Ron Kimmel for useful discussions. Some figures in this paper were created using BioRender [27].

## 6 Code Availability

The code to reproduce the figures in this paper is available at https://github.com/yevgenin/DeepOM.

## 7 Data Availability

The data underlying this paper will be shared upon reasonable request to the corresponding author.

## References

[1] Michal Levy-Sakin and Yuval Ebenstein. Beyond sequencing: optical mapping of dna in the age of nanotechnology and nanoscopy. Current opinion in biotechnology, 24(4):690–698, 2013.

[2] Vilhelm Müller and Fredrik Westerlund. Optical dna mapping in nanofluidic devices: principles and applications. Lab on a Chip, 17(4):579–590, 2017.

[3] Dominika Gruszka, Jonathan Jeffet, Sapir Margalit, Yael Michaeli, and Yuval Ebenstein. Single-molecule optical genome mapping in nanochannels: multidisciplinarity at the nanoscale. Essays in Biochemistry, 65(1):51–66, 2021.

[4] Arno Bouwens, Jochem Deen, Raffaele Vitale, Laurens D’Huys, Vince Goyvaerts, Adrien Descloux, Doortje Borrenberghs, Kristin Grussmayer, Tomas Lukes, Rafael Camacho, Jia Su, Cyril Ruckebusch, Theo Lasser, Dimitri Van De Ville, Johan Hofkens, Aleksandra Radenovic, and Kris Pieter Frans Janssen. Identifying microbial species by single-molecule dna optical mapping and resampling statistics. NAR Genomics and Bioinformatics, 2:1–13, 3 2020.

[5] Nathaniel O Wand, Darren A Smith, Andrew A Wilkinson, Ashleigh E Rushton, Stephen J W Busby, Iain B Styles, and Robert K Neely. Dna barcodes for rapid, whole genome, single-molecule analyses. Nucleic Acids Research, 47:e68–e68, 7 2019.

[6] Assaf Grunwald, Moran Dahan, Anna Giesbertz, Adam Nilsson, Lena K. Nyberg, Elmar Weinhold, Tobias Ambjörnsson, Fredrik Westerlund, and Yuval Ebenstein. Bacteriophage strain typing by rapid single molecule analysis. Nucleic Acids Research, 43, 10 2015.

[7] Vilhelm Müller, My Nyblom, Anna Johnning, Marie Wrande, Albertas Dvirnas, Sriram KK, Christian G. Giske, Tobias Ambjörnsson, Linus Sandegren, Erik Kristiansson, and Fredrik Westerlund. Cultivation-free typing of bacteria using optical dna mapping. ACS Infectious Diseases, 6:1076–1084, 5 2020.

[8] Dmitry Torchinsky, Yael Michaeli, Natalie R Gassman, and Yuval Ebenstein. Simultaneous detection of multiple dna damage types by multi-colour fluorescent labelling. Chemical Communications, 55(76):11414–11417, 2019.

[9] Hila Sharim, Assaf Grunwald, Tslil Gabrieli, Yael Michaeli, Sapir Margalit, Dmitry Torchinsky, Rani Arielly, Gil Nifker, Matyas Juhasz, Felix Gularek, Miguel Almalvez, Brandon Dufault, Sreetama Sen Chandra, Alexander Liu, Surajit Bhattacharya, Yi-Wen Chen, Eric Vilain, Kathryn R. Wagner, Jonathan Pevsner, Jeff Reifenberger, Ernest T. Lam, Alex R. Hastie, Han Cao, Hayk Barseghyan, Elmar Weinhold, and Yuval Ebenstein. Long-read single-molecule maps of the functional methylome. Genome Research, 29:646–656, 4 2019.

[10] Peter Ebert, Peter A Audano, Qihui Zhu, Bernardo Rodriguez-Martin, David Porubsky, Marc Jan Bonder, Arvis Sulovari, Jana Ebler, Weichen Zhou, Rebecca Serra Mari, et al. Haplotype-resolved diverse human genomes and integrated analysis of structural variation. Science, 372(6537):eabf7117, 2021.

[11] Sapir Margalit, Yotam Abramson, Hila Sharim, Zohar Manber, Surajit Bhattacharya, Yi-Wen Chen, Eric Vilain, Hayk Barseghyan, Ran Elkon, Roded Sharan, et al. Long reads capture simultaneous enhancer–promoter methylation status for cell-type deconvolution. Bioinformatics, 37(Supplement 1):i327–i333, 2021.

[12] Mickaël Lelek, Melina T Gyparaki, Gerti Beliu, Florian Schueder, Juliette Griffié, Suliana Manley, Ralf Jungmann, Markus Sauer, Melike Lakadamyali, and Christophe Zimmer. Single-molecule localization microscopy. Nature Reviews Methods Primers, 1(1):1–27, 2021.

[13] Anton Valouev, Lei Li, Yu-Chi Liu, David C Schwartz, Yi Yang, Yu Zhang, and Michael S Waterman. Alignment of optical maps. Journal of Computational Biology, 13(2):442–462, 2006.

[14] Sapir Margalit, Zuzana Tulpova, Yael Michaeli, Tahir Detinis Zur, Jasline Deek, Sivan Louzoun-Zada, Assaf Grunwald, Yuval Scher, Leonie Schutz, Elmar Weinhold, et al. Optical genome and epigenome mapping of clear cell renal cell carcinoma. bioRxiv, 2022.

[15] Tslil Gabrieli, Hila Sharim, Gil Nifker, Jonathan Jeffet, Tamar Shahal, Rani Arielly, Michal Levi-Sakin, Lily Hoch, Nissim Arbib, Yael Michaeli, et al. Epigenetic optical mapping of 5-hydroxymethylcytosine in nanochannel arrays. ACS nano, 12(7):7148–7158, 2018.

[16] Tslil Gabrieli, Yael Michaeli, Sigal Avraham, Dmitry Torchinsky, Sapir Margalit, Leonie Schütz, Matyas Juhasz, Ceyda Coruh, Nissim Arbib, Zhaohui Sunny Zhou, et al. Chemoenzymatic labeling of dna methylation patterns for single-molecule epigenetic mapping. Nucleic acids research, 50(16):e92–e92, 2022.

[17] Elias Nehme, Lucien E Weiss, Tomer Michaeli, and Yoav Shechtman. Deep-storm: super-resolution single-molecule microscopy by deep learning. Optica, 5(4):458–464, 2018.

[18] Elias Nehme, Daniel Freedman, Racheli Gordon, Boris Ferdman, Lucien E Weiss, Onit Alalouf, Tal Naor, Reut Orange, Tomer Michaeli, and Yoav Shechtman. Deepstorm3d: dense 3d localization microscopy and psf design by deep learning. Nature methods, 17(7):734–740, 2020.

[19] Artur Speiser, Lucas-Raphael Müller, Philipp Hoess, Ulf Matti, Christopher J Obara, Wesley R Legant, Anna Kreshuk, Jakob H Macke, Jonas Ries, and Srinivas C Turaga. Deep learning enables fast and dense single-molecule localization with high accuracy. Nature methods, 18(9):1082–1090, 2021.

[20] Thomas Anantharaman and Bud Mishra. False positives in genomic map assembly and sequence validation, 2001.

[21] Fausto Milletari, Nassir Navab, and Seyed-Ahmad Ahmadi. V-net: Fully convolutional neural networks for volumetric medical image segmentation. In 2016 fourth international conference on 3D vision (3DV), pages 565–571. IEEE, 2016.

[22] Carole H Sudre, Wenqi Li, Tom Vercauteren, Sebastien Ourselin, and M Jorge Cardoso. Generalised dice overlap as a deep learning loss function for highly unbalanced segmentations. In Deep learning in medical image analysis and multimodal learning for clinical decision support, pages 240–248. Springer, 2017.

[23] Charles J Clopper and Egon S Pearson. The use of confidence or fiducial limits illustrated in the case of the binomial. Biometrika, 26(4):404–413, 1934.

[24] Sergey Nurk, Sergey Koren, Arang Rhie, Mikko Rautiainen, Andrey V Bzikadze, Alla Mikheenko, Mitchell R Vollger, Nicolas Altemose, Lev Uralsky, Ariel Gershman, et al. The complete sequence of a human genome. Science, 376(6588):44–53, 2022.

[25] Bionano documentation. https://bionanogenomics.com/support-page/data-analysis-documentation/. Accessed: 2022-10-19.

[26] Rani Arielly and Yuval Ebenstein. Irys Extract. Bioinformatics, 34(1):134–136, 07 2017.

[27] Some figures in this paper were created with biorender.com.

